## Supplemental Analysis, Figures, Tables, Dataset Captions for "Global pathogenomic analysis identifies known and novel genetic antimicrobial resistance determinants in twelve species"

### Supplemental Analysis, Figures, Tables, and Dataset Captions

#### Supplemental Analysis

*Diversity of selected genomes*

The overall diversity of the genomes selected for this study was evaluated by examining the distribution of MLST subtypes (from https://github.com/tseemann/mlst) and BioProject IDs (from PATRIC) for each genome, and the number of susceptible/resistant genomes per drug (Figure S1, Dataset S2). MLST distributions suggest that the genome collection for each species is genetically diverse, with the most common subtype per species never exceeding 50% of all genomes and at least 44 subtypes represented per species. BioProject ID distributions suggest that a majority of our genome collections also represent a wide range of studies, though the smallest collections were dominated by individual studies (*C. coli* and *C. jejuni* genomes originated almost entirely from PRJNA292668, *E. cloacae* genomes were majority from PRJEB5065) and genomes for *E. faecium* and *N. gonorrhoeae* had poor coverage regarding study of origin. Finally, genome counts by resistance suggest that both susceptible and resistant strains across a wide range of drug classes are present for most species, with only the *N. gonorrhoeae* collection potentially being limited due to AMR data being available for only two drugs.

*Observation of TEM-family beta-lactamases in both gram-positive and gram-negative strains*

Given the prevalence of TEM family beta-lactamases (blaTEMs) among the few AMR genes spanning multiple phylogenetic classes, all observed blaTEM alleles were mapped to known blaTEM alleles based on RGI annotations, focusing on the 51 “complete” alleles with length within 5% of the TEM-1 allele. Complete blaTEMs were observed in 4,861 genomes spanning 8 species and were dominated by the TEM-1 allele occurring in 4,424 genomes (Figure S4a, Table S3, Dataset S4). All but five alleles were within two mutations of TEM-1, and only 9 alleles (including TEM-1) were observed in at least 10 genomes. Individual alleles were largely specific to phylogenetic class with 38 alleles limited to Gammaproteobacteria and 10 alleles limited to *N. gonorrhoeae*, compared to two alleles observed in both (TEM-1 and TEM-

135). Just one allele was observed in gram-positive strains, TEM-116 (substitutions V82I and A182V relative to TEM-1), occurring in 11 *S. aureus* strains and 17 *S. enterica* strains. Contigs from genome assemblies harboring TEM-116 were mapped to known plasmids on PLSDB[1] using MASH[2], and the 28 instances of TEM-116 were predicted to be located on one of three plasmids: NZ_AJ437107.1 (14 *S. enterica*, 1 *S. aureus*), NZ_AJ438270.1 (3. *S. enterica*), and NC_019053.1 (10 *S. aureus*) (Figure S4b-c, Dataset S4).

Pairwise MASH distances between these genomes were compared to those between all *S. aureus* and *S. enterica* genomes to assess sequence similarity and clonality (Figure S5). The 17 *S. enterica* genomes with TEM-116 were highly similar with a median pairwise MASH distance of 0.0004 compared to a median of 0.0153 between all *S. enterica* genome pairs, consistent with the source study confirming all such genomes to be derived from a single clonal lineage of serotype Kentucky ST198, albeit from geographically diverse locations[3]. However, as the presence of the two TEM-116 plasmids in these genomes do not closely track their phylogeny, it is ambiguous as to whether the observed plasmid distribution was established earlier in the lineage and resulted from clonal dissemination, or was a result of more recent horizontal gene transfer events.

In contrast, the 11 *S. aureus* strains with TEM-116 were genetically diverse with a median pairwise MASH distance of 0.0080 compared to a median of 0.0151 between all *S. aureus* genome pairs. The genome 1280.16776, which is predicted to carry plasmid NZ_AJ437107.1 shared by some *S. enterica* genomes, is genetically distinct from the other TEM-116 carrying *S. aureus* genomes. Given the geographic prevalence of the *S. enterica* genomes carrying NZ_AJ437107.1 (annotated as isolated across 10 countries: Djibouti, Egypt, France, Indonesia, Israel, Kenya, Kuwait, Morocco, Tanzania, Vietnam)[3], it is possible that the strain corresponding to 1280.16776 could have acquired this plasmid via horizontal gene transfer from a gram-negative species.

To assess the impact of blaTEM on beta-lactam resistance in *S. aureus*, the presence of blaTEM and other beta-lactam AMR genes was compared to cefoxitin MIC data, which was available for *S. aureus* genomes harboring blaTEM. While only four genomes with blaTEM also had MIC data, those genomes (which also carried *mecA* and *blaZ*) had significantly elevated MICs compared to those with just *mecA* and *blaZ* or *mecA* alone (p=0.0074, Mann-Whitney U-test), suggesting blaTEM may confer additional protection from beta-lactams in already resistant *S. aureus* (Figure S4c, Figure S6). These four strains were isolated from a study of the pork production chain in Shandong, China (BioProject: PRJNA433074)[4], reflecting a geographically confined but genetically diverse sample set. The strains span two MLST subtypes (ST9, ST59), with three isolated from workers (PATRIC IDs: 1280.16838, 1280.16865, 1280.16862) and one from a pig (1280.16776).

*Association between features annotated as known AMR genes and resistance*

For each known AMR feature across the 127 species-drug cases examined using the SVM workflow, the set of genomes carrying the feature was identified and the fraction of such genomes that are resistant to the corresponding drug was computed. The distribution of these resistance fractions was computed for each species-drug case individually and in aggregate, combining all known AMR features across all cases (Figure S14). The distributions generally concentrated around resistance fractions of 0 or 1, which was quantified by computed the mean of resistance fractions for values either less than or greater than 0.5, for each species-drug case (Figure S14b-c). These results suggest that features labeled as "known AMR" are more likely than not related to resistance and are correctly labeled, i.e. features with resistance fractions closer to 1 are more likely to be resistance determinants, while those with fractions closer to 0 are more likely to be markers for susceptibility such as known fluoroquinolone-susceptible *gyrA* alleles.

*Design of the GWAS score for biological relevance*

The "GWAS score" was inspired by some of the informal intuition behind how follow-up analyses are often conducted for the top hits from an association study. When hundreds of significant hits are observed, many GWAS and ML studies of AMR will focus on the top X features by effect size (i.e. top 3, 5, 10, 20)[5-7]. The GWAS score mimics this diminishing attention with respect to rank by giving exponentially diminishing marginal GWAS score per recovered known gene as rank increases, specifically, to diminish by half every 10 ranks.

*Support vector machine hyperparameter assessment on test cases*

Ten test species-drug cases were selected among those with substantial data (at least 1000 SIRs and 50 known AMR genes), with balanced representation by drug class and species phylogenetic class (Table S4). Across these cases, we find that ensemble size had little impact on either accuracy or biological relevance, while the other HPs significantly impacted both metrics: SVM regularization term C, fraction of samples per estimator, and fraction of features per estimator (Kruskal-Wallis test, FWER < 0.05, Bonferroni correction, 80 tests) (Figure S7, Figure S8a, Table S5). More aggressive feature subsampling (smaller fraction of features per estimator) also consistently improved on biological relevance without compromising accuracy (Figure S8b). Applying Mann-Whitney U-tests to test for differences in GWAS scores from using feature fraction 25% over 50%, 50% over 75%, and 75% over 100% yielded p-values of $3.0*10^{-17}$, $6.0*10^{-38}$, and $2.6*10^{-29}$, respectively. Analogous tests for differences in test set MCC yielded $1.5*10^{-7}$, 0.03, and 0.27, respectively, of which only the test between 25% and 50% is significant at FWER < 0.05 (Bonferroni correction, 6 tests). Finally, the smallest subset of HP combinations containing nearly-optimal models (within 90% of maximum MCC and GWAS scores) was identified for more efficient HP optimization of the remaining species-drug cases (Table S6).

The impact of reducing the number of tested HP combinations was examined as follows. For each of the 10 test species-drug cases, the test set MCCs and GWAS scores across all folds (from 5-fold cross validation) were combined across all models resulting from tested HP combinations, then split into those associated with included HP combinations and those associated with excluded combinations. Mann-Whitney U-tests were applied to determine whether the MCCs or GWAS scores differed significantly between models with included or excluded HP combinations (Table S7). Significant differences were observed in 2/10 cases for MCCs and 5/10 cases for GWAS scores (FWER < 0.05, Bonferroni correction, 20 tests). In 2/2 significant MCC cases and 3/5 significant GWAS score cases, the subset had an equal or higher median score than the excluded combinations, suggesting that the reduction in the set of tested HP combinations only rarely reduces the maximum performance attainable.

*Variations on the preliminary feature filter*

Variations to the approach of pre-filtering features by log odds ratio (LOR) were evaluated for their impact on downstream model performance. The existing filter of taking the top 50,000 features by LOR (specifically, the features with the 25,000 highest and lowest LORs) was compared to analogous filters taking the top 20,000, 10,000, or 5,000 features by LOR. These were also compared to filters taking the top 50,000, 20,000, 10,000 or 5,000 features by Fisher's exact test p-value when testing for enrichment in resistant genomes, for a total of eight possible feature filters. After applying each filter, SVM ensembles were trained to predict AMR phenotype across the 10 test species-drug cases and 256 HP combinations described in the previous section. The maximum and median test set MCC (from 5-fold cross validation) and GWAS score was computed for each species-drug case and under each filter (Figure S15, Dataset S5).

In most of the tested species-drug cases, the maximum and median MCC and GWAS scores are robust to the number of features provided, with a few exceptions under Fisher's exact test filters: Model MCCs for the *E. coli*-gentamicin and *S. aureus*-erythromycin cases and GWAS scores for the *A. baumannii*-amikacin, *S. aureus*-ciprofloxacin, and *S. aureus*-erythromycin cases increase with the number of features. When comparing equal size feature sets generated from filtering by either LOR or Fisher's exact test p-value, models derived from the LOR filter consistently performed better. These results suggest that 1) future applications of this ML workflow may be able to apply stricter preliminary feature filters to reduce the computational resources required for training with relatively little impact on performance, and 2) the LOR appears to more effectively identify features required for better performing models than the Fisher's exact test p-value, possibly due to its emphasis on effect size over significance.

*Generalizability of the GWAS score*

To assess whether a model's GWAS score calculated from a fixed set of known AMR genes is representative of its capacity to recover other "unseen" AMR genes, the following experiment was conducted. For a given species-drug case, 1) half of all known AMR genes were randomly hidden, 2) GWAS scores were computed using either visible or hidden AMR genes across all 256 tested hyperparameter combinations, and 3) the Spearman correlation was computed between the two GWAS scores as a measure of generalizability. This experiment was conducted for 100 random selections of hidden AMR genes for each of the 10 test species-drug cases (Dataset S5). Overall, the results suggest that the GWAS score often generalizes well to hidden AMR genes, with a median Spearman correlation between GWAS scores from visible vs. hidden AMR genes exceeding 0.4 in 6/10 species-drug cases and 0.6 in 4/10 cases (Figure S9a). The extent of generalizability was dependent on the initial level of GWAS score variation (computed as the standard deviation of GWAS scores across all hyperparameter combinations with all AMR genes visible) (Figure S9b). Low initial variation in GWAS scores may result in a less robust ranking of models by GWAS score, and consequently weaker correlations between GWAS scores calculated using different subsets of known AMR genes. Conversely, cases of high initial variation in GWAS score, i.e. those where HP optimization can meaningfully impact model performance, are also more likely to have GWAS scores that are representative of the model's performance at recovering yet unknown AMR genes.

*Effect of dataset parameters on model performance*

The relationship between six dataset parameters and model performance was analyzed: species, drug class, dataset size (number of genomes), extent of class imbalance (minority phenotype fraction), fraction of genomes originally assigned the "intermediate" phenotype, and total number of known AMR genes annotated (Figure S10, Table S8). Associations between these parameters and either the model's predictive performance (mean test set MCC from 5-fold cross validation) or known AMR gene recovery (number of AMR genes recovered among the top 20 features) were examined using Spearman R tests for quantitative parameters and Kruskal-Wallis tests for qualitative parameters. Of these, four tests were significant at FWER < 0.05 (Bonferroni correction, 12 tests): Number of genomes vs. MCC, species vs. MCC, fraction of intermediate genomes vs. AMR gene recovery, and total known AMR genes vs. AMR gene recovery.

The significant result regarding intermediate genomes points to an opportunity to improve the overall workflow. Given the negative association between intermediate genomes and model performance (likely due to their genetic similarity to both susceptible and resistant genomes), future iterations of this workflow may benefit from treating intermediate genomes as a separate phenotype, and original SIR and MIC phenotypes are available in Dataset S2. Nonetheless, the prevalence of intermediate genomes and their potential confounding effects is unlikely to have a significant impact on the final set of AMR gene candidates, as the accurate models (those with MCC > 0.8) used for identifying candidates had a median intermediate genome fraction of only 0.8%. Finally, we note the weak but negative correlation between the number of known AMR genes and model MCC (Spearman correlation = -0.215). This result is consistent with previous ML studies of AMR suggesting that models that accurately predict AMR phenotype often do not rely on known AMR genes and highlights the importance of recognizing phenotype prediction accuracy and recovery of true AMR genes as distinct objectives.

*Characterization of gyrA alleles associated with fluoroquinolone resistance recovered using support vector machine ensembles*

All *gyrA* alleles among the top 50 features from models related to fluoroquinolone resistance were identified, totaling 30 alleles across 7 species. Mutations were called relative to the wildtype allele of the corresponding species, defined as the most commonly observed *gyrA* allele among genomes susceptible to at least one fluoroquinolone. Resulting mutations and GenBank accession IDs for wildtype alleles are available in Dataset S5. All recovered *gyrA* alleles with positive feature weight for resistance had at least one known resistance-conferring mutation, covering the following substitutions: S81L in *A. baumannii*[8], T86I in *C. coli* and *C. jejuni*[9], S83L and D87N in *E. coli*[10], T83I in *P. aeruginosa*[11], and S83Y and D87* in *S. enterica*[12]. All recovered *gyrA* alleles with negative feature weight for resistance had no such known resistance-conferring mutations. These results suggest that the *gyrA* alleles recovered by the SVM ensemble approach are consistent with current understanding of the *gyrA* mutational landscape with respect to fluoroquinolone resistance.

*Selection of AMR gene candidates for experimental validation*

AMR gene candidates were identified through a series of filters applied to the most predictive features in the most accurate AMR models (Figure 4a). The top 10 features by weight across the 78 species-drug AMR models achieving test MCC > 0.8 were combined to yield an initial set of 886 unique genetic features or "AMR-predictive genetic features". Of these, 610 were not directly associated with known AMR genes, 519 were also observed in at least 10 genomes, and 347 also had positive log odds ratios (LORs) for resistance against the drug for which the feature was predictive of resistance. Candidates in this intermediate set were then scored based on the sum of the following: 1) number of drugs in the same drug class for which the feature is significantly associated with resistance (applying Fisher's exact test to SIR data or applying Brunner-Munzel test to MIC data), and 2) number of drugs in the same drug class for which the feature appears in at least one resistant genome without any other known AMR genes (i.e. number of drugs for which this feature could explain previously unexplainable resistance). Finally, for each species-drug class pair, the top 10 features by this score were selected to yield the 142 AMR gene candidates (some pairs had fewer than 10 features remaining after the initial set of filters). Results of the scoring process are detailed in Dataset S7.

In the selection of features for experimental validation, the 142 candidates were further categorized by function and filtered down to 43 features that were 1) not poorly characterized, 2) not associated with mobile elements (transposases, insertion elements, phage elements, integrases, plasmid maintenance), 3) not AMR genes for unrelated drugs, and 4) not strongly correlated with a known AMR gene after manual inspection (Figure 4b). Details regarding these 43 features are in Dataset S7. Of these, six features referred to specific sequence variants in *E. coli,* the species the authors were best equipped to test experimentally. Two were excluded in order to focus on allele-type features and take advantage of the Keio knockout collection[13], and ultimately *frdD* and *cycA* were selected for experimental validation for their better functional and metabolic characterization compared to the other two options, *sugE* and *yjfN*.

*Distribution of cycA, frdD, and ampC across E. coli genomes*

Analysis of all 3,856 *E. coli* genomes in this study suggests that *cycA*, *frdD*, and *ampC* are core genes of *E. coli*, found in 3,823 (99.1%), 3,796 (98.4%), and 3,775 (97.8%) of genomes, respectively. No genomes had multiple copies of *cycA*, *frdD*, or *ampC*. 3,748 (97.2%) genomes had both *frdD* and *ampC*, of which all but three harbored *frdD* and *ampC* on the same contig. The distance between the *frdD* and *ampC* ORFs was highly consistent with a mean of 62.9bp, standard deviation of 5.2bp, and range of 30-192bp. A vast majority of genomes (3,698) had a *frdD*-*ampC* distance of exactly 63bp. This result suggests that a *frdD* V111D mutation will impact the *ampC* promoter in most *E. coli* strains. All instances of *cycA*, *frdD*, and *ampC* identified across all *E. coli* genomes are available in Dataset S8.

### Supplemental Analysis – References

### Supplemental Figures

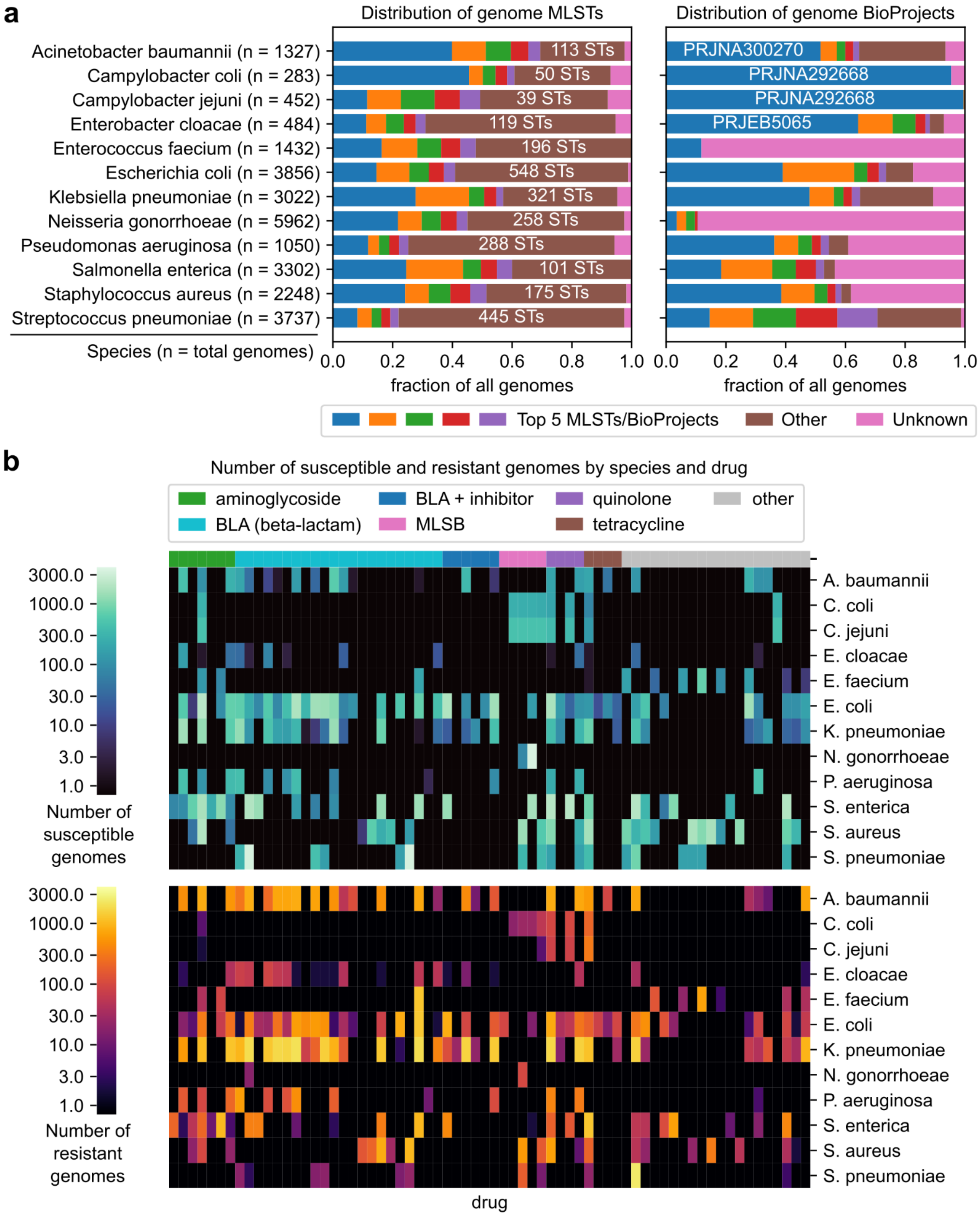


**Figure S1: Diversity of samples by subtype, study of origin, and AMR phenotype.** a) Distribution of MLST subtypes and BioProject accession IDs for each species' genome collection. For MLSTs, the total number of additional MLST subtypes beyond the five most common are labeled. BioProjects comprising at least 50% of genomes for a given species are labeled. b) Distribution of AMR phenotypes by species, drug, and drug class. Drugs are sorted by drug class. The macrolide-lincosamide-streptogramin B drug class is abbreviated “MLSB”.

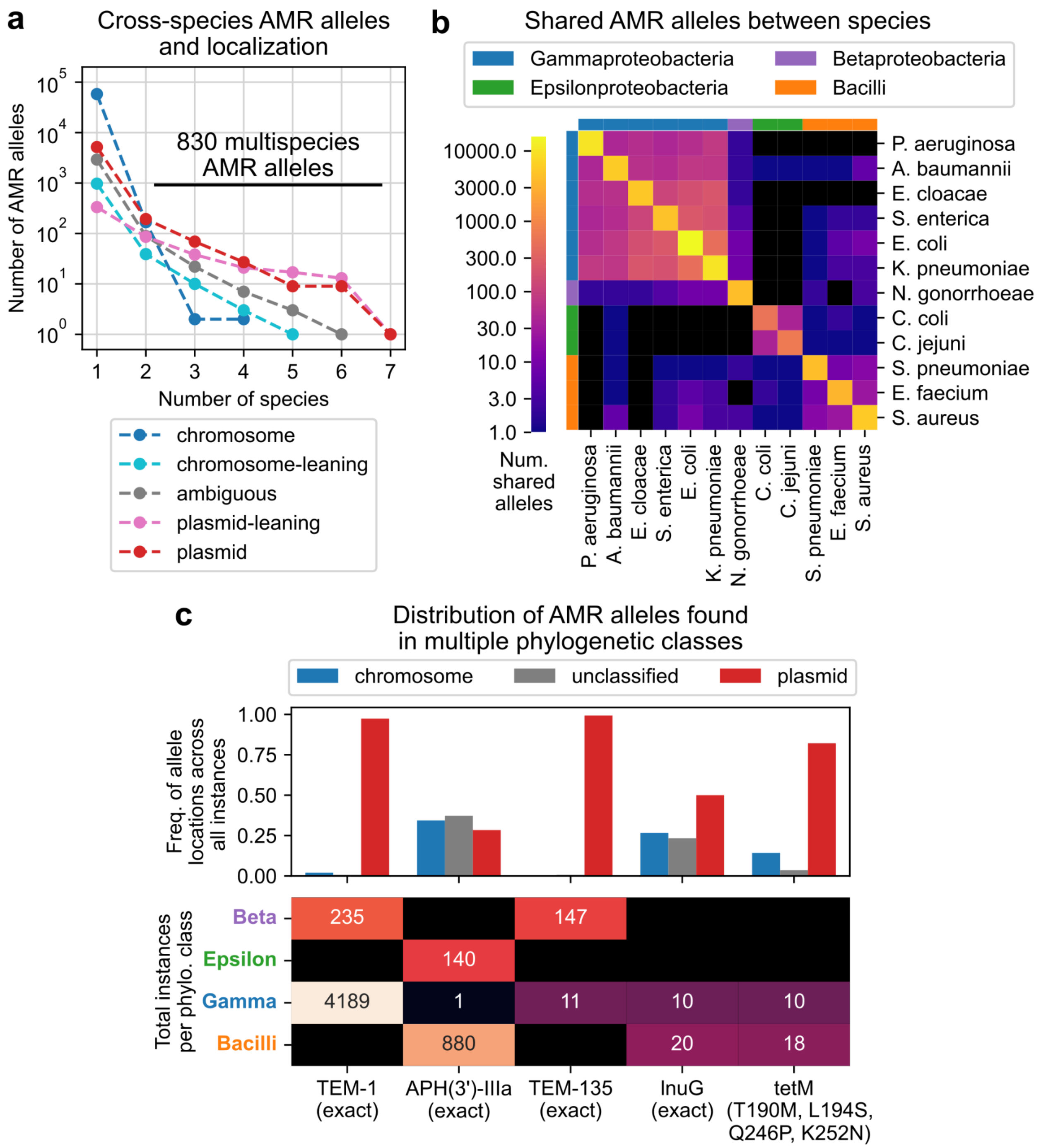


**Figure S2: Cross-species analysis of 68,324 antimicrobial resistance gene alleles and gene locations.** a) Relationship between the number of species an AMR allele is observed in and tendency to be plasmid-encoded. b) Number of AMR alleles shared between each pair of species, compared to species phylogenetic class. d) Distribution of predicted gene locations and total occurrences per phylogenetic class for AMR alleles appearing in at least 10 genomes in multiple classes.

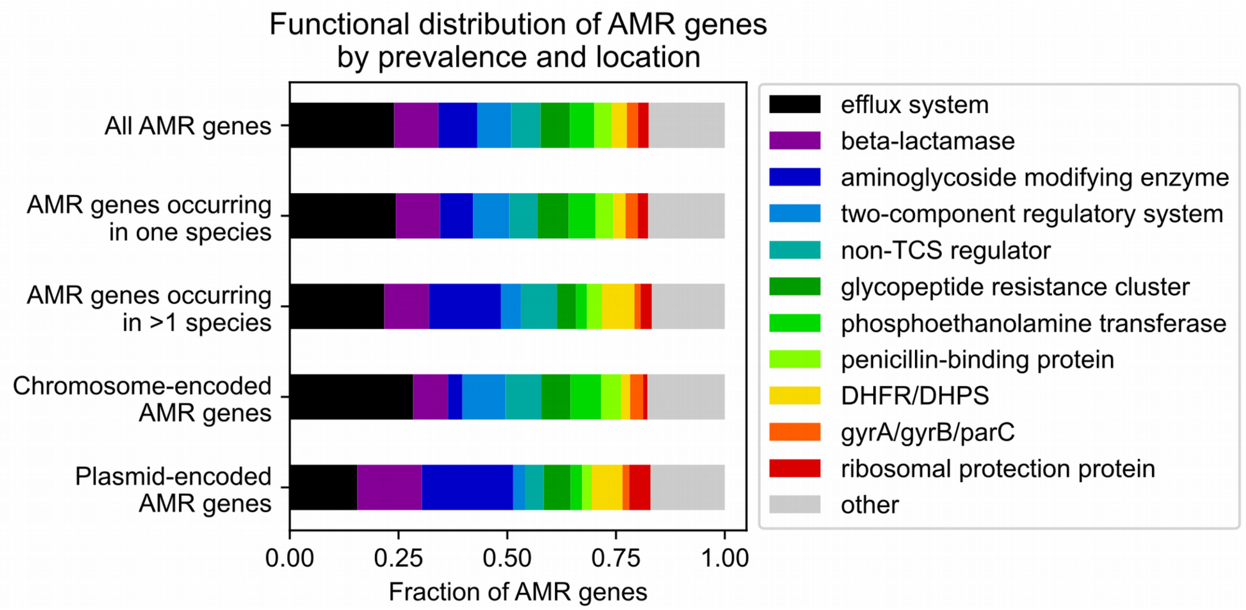


**Figure S3: Distribution of 6,332 antimicrobial resistance genes across 12 species by functional category, cross-species prevalence, and gene location.** Abbreviated functions are two-component regulatory system (TCS), dihydrofolate reductase (DHFR), and dihydropteroate synthase (DHPS).

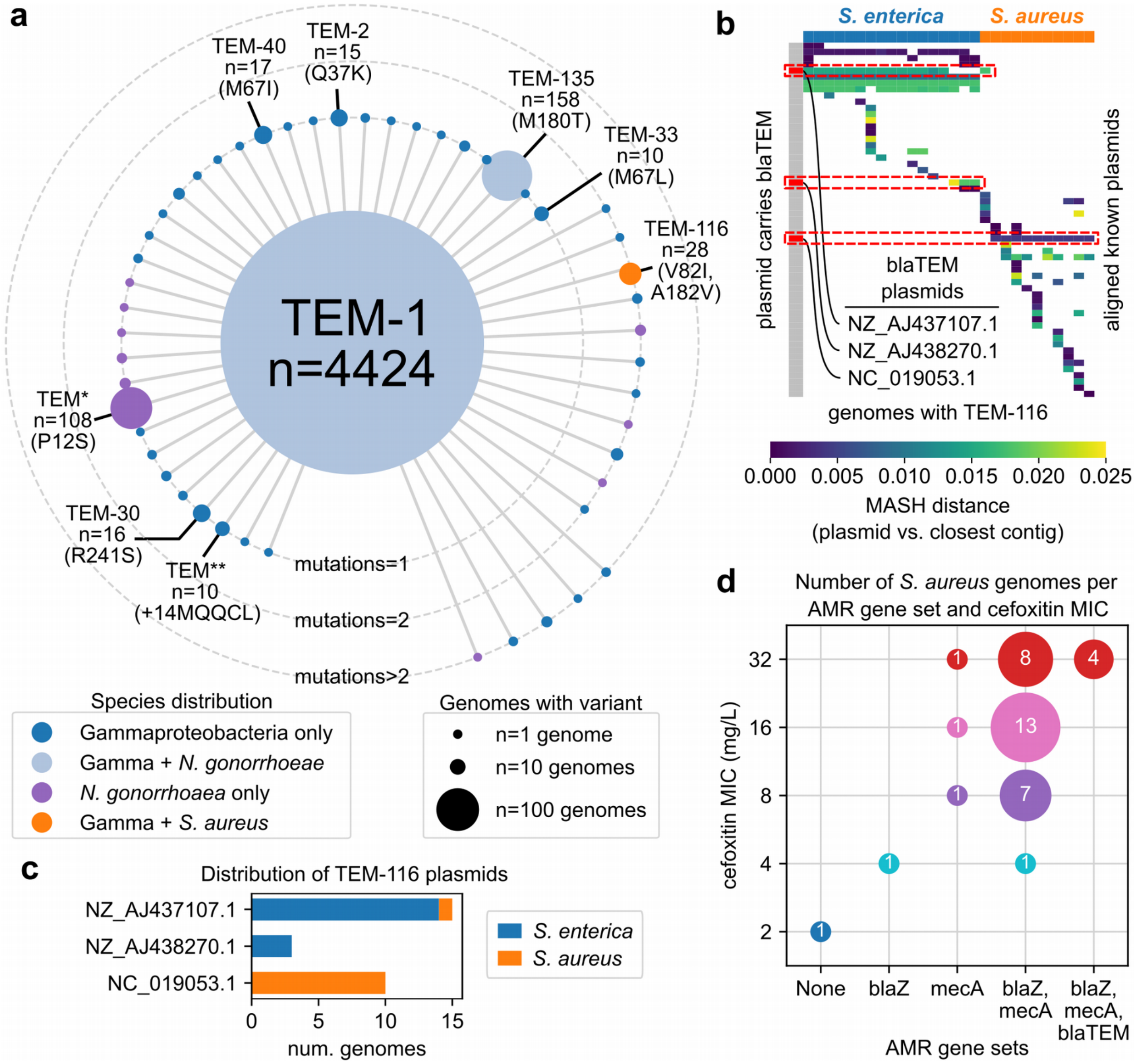


**Figure S4: Distribution of TEM-family beta-lactamases observed in 4,861 genomes across 8 species.** a) Distribution of TEM-family beta-lactamase (blaTEM) alleles with respect to genome count, harboring species, and mutations relative to the TEM-1 allele. Alleles occurring in at least 10 genomes are labeled. TEM* and TEM** refer to unnamed blaTEM alleles. b) Alignments between contigs in genomes with TEM-116 and PLSDB plasmids based on MASH distance. Known plasmids carrying blaTEM are indicated. All plasmids with MASH distance < 0.025 to at least one genome with TEM-116 are shown. c) Distribution of the three plasmids containing TEM-116 by species. d) Relationship between beta-lactam associated AMR genes and cefoxitin minimum inhibitory concentration (MIC) in *S. aureus*. Numeric labels correspond to the number of genomes observed with the corresponding AMR genes and MIC.

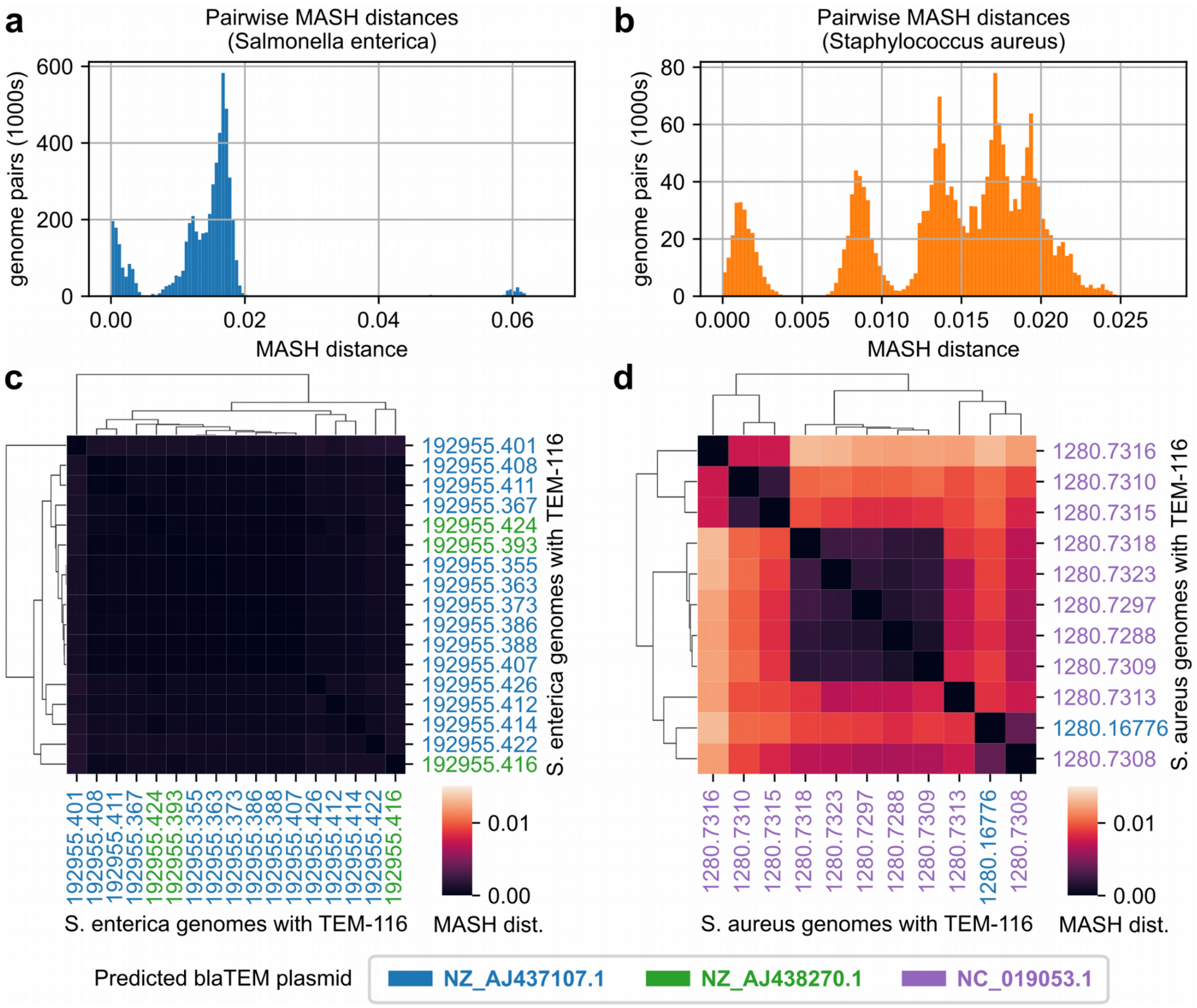


**Figure S5: Sequence similarity between *S. enterica* and *S. aureus* genomes carrying TEM-116.** a-b) Distributions of pairwise MASH distances between all 3,302 *S. enterica* and 2,248 *S. aureus* genomes. c-d) Clustermaps based on pairwise MASH distances between the 17 *S. enterica* and 11 *S. aureus* genomes carrying TEM-116. Heatmaps share the same color scales. Genomes are colored by their predicted blaTEM plasmid and are clustered using single linkage and Euclidean distances.

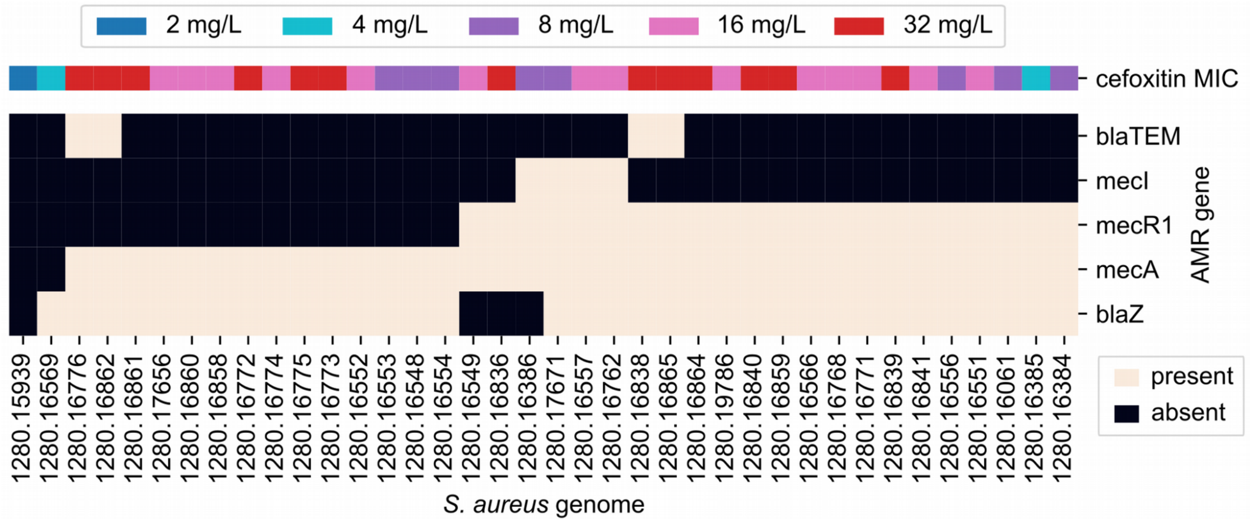


**Figure S6: Cefoxitin MIC versus beta-lactam resistance-associated genes in *S. aureus*.** Rows and columns have been ordered by hierarchical clustering with Jaccard distances and average linkage.

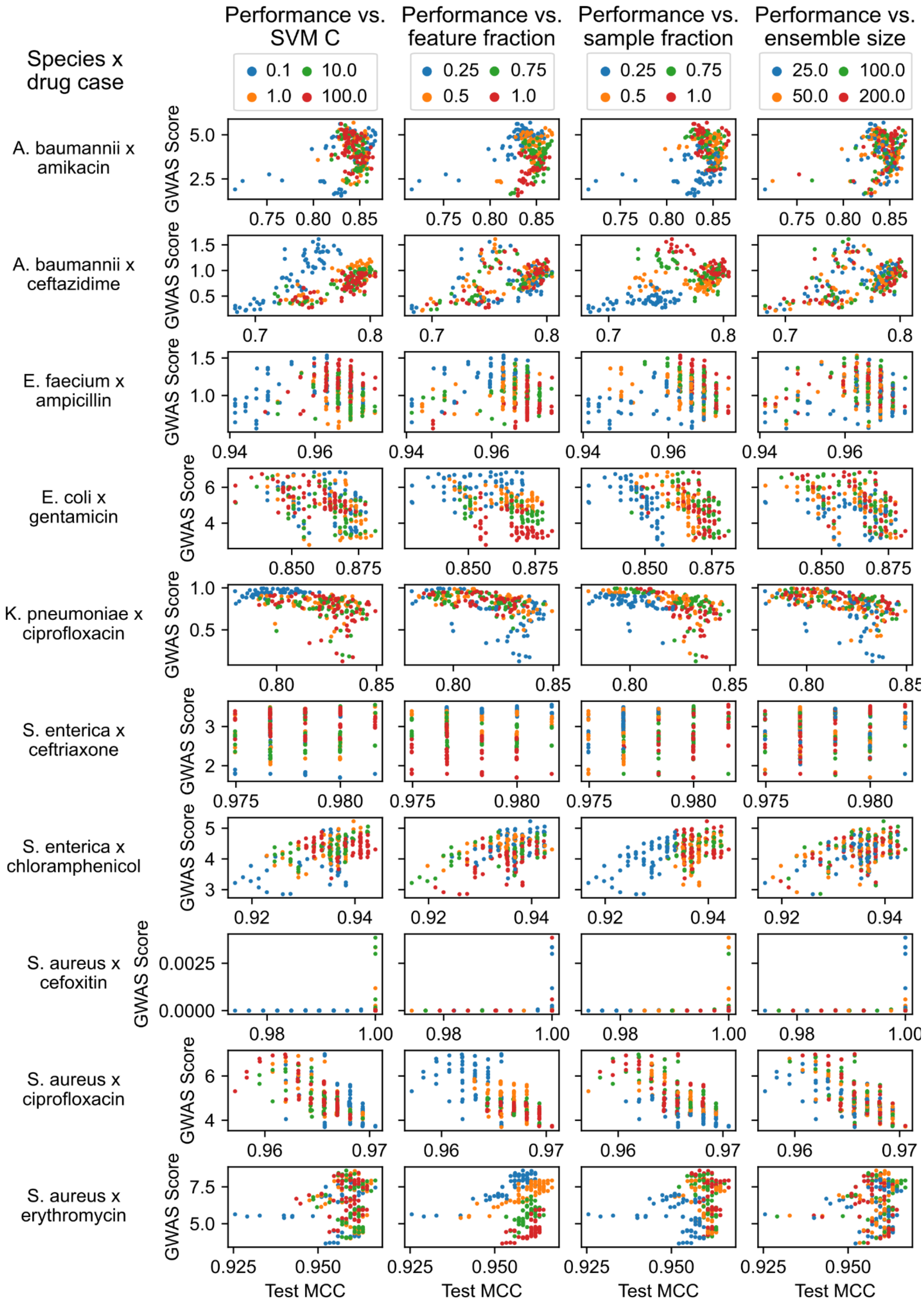


**Figure S7: Impact of SVM ensemble hyperparameters on AMR phenotype prediction performance and recovery of known AMR genes.** Each row corresponds to a single AMR prediction problem between a species and a drug, and each column corresponds to a varied hyperparameter. X-axes show average MCCs on the test set from 5-fold cross validation (CV) experiments, y-axes show average GWAS scores across the CV experiment.

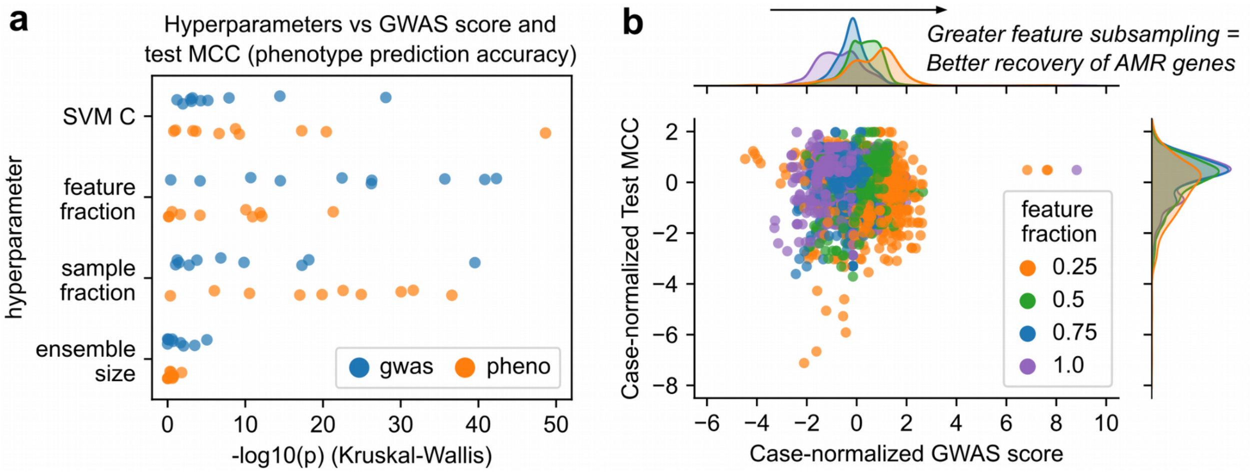


**Figure S8: Overall impact of SVM ensemble hyperparameters on performance.** a) Kruskal-Wallis ANOVA tests between hyperparameters and either AMR phenotype prediction performance (Matthew's correlation coefficient; MCC) or biological relevance (GWAS score) across 10 test species-drug cases. b) Impact of feature subsampling on model performance for the 10 test cases. MCCs and GWAS scores have been normalized to mean 0 and standard deviation 1, within their specific species-drug cases.

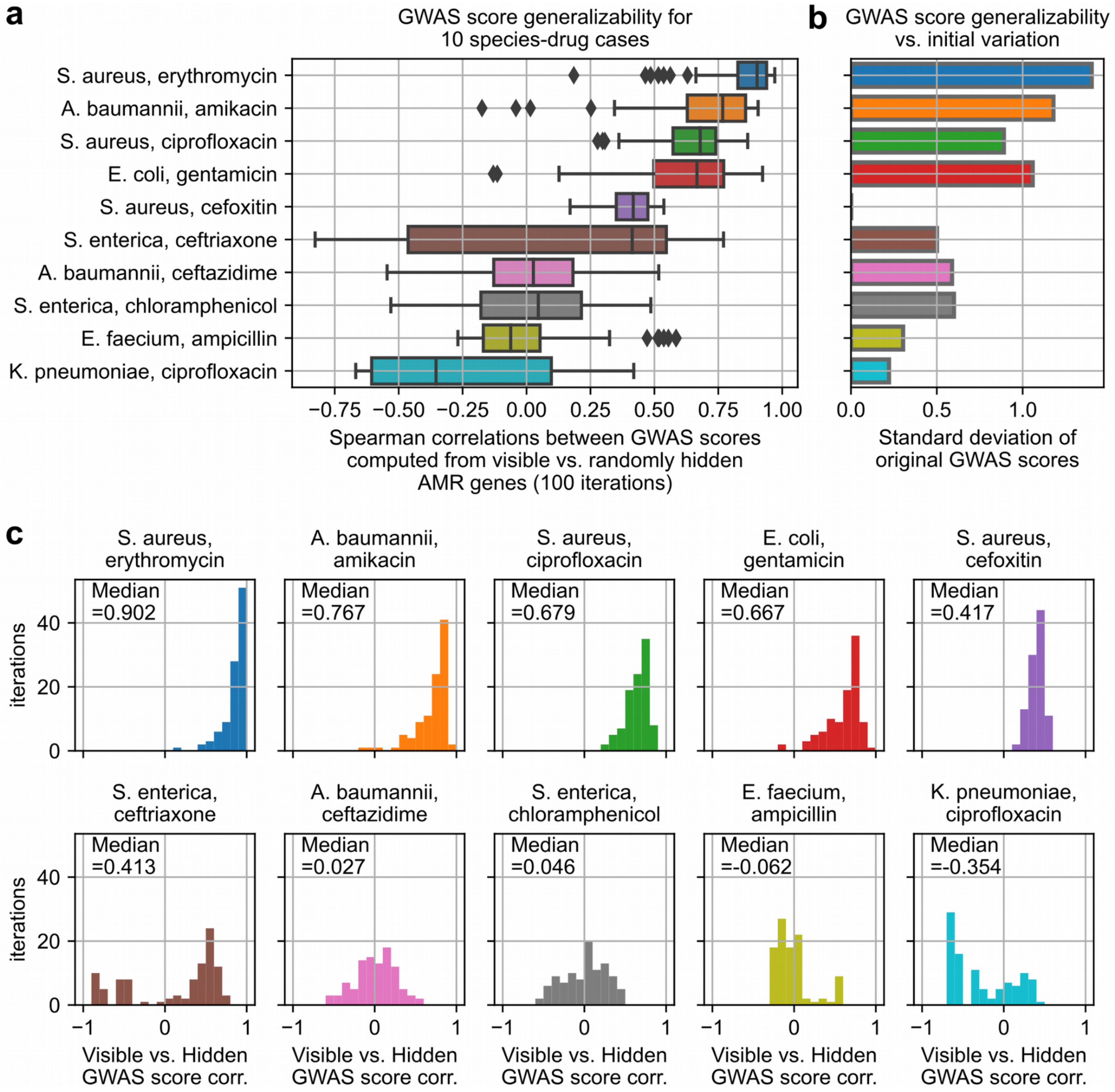


**Figure S9: Generalizability of the GWAS score.** For each species-drug case, half of all known AMR genes were hidden randomly and GWAS scores were computed using either visible or hidden AMR genes for models trained using each possible hyperparameter combination. The distribution of Spearman correlations between the two GWAS scores is shown in (a) for 100 random selections of hidden AMR genes. b) Standard deviation of GWAS scores per species-drug case. c) Distribution of GWAS score Spearman correlations by species-drug case.

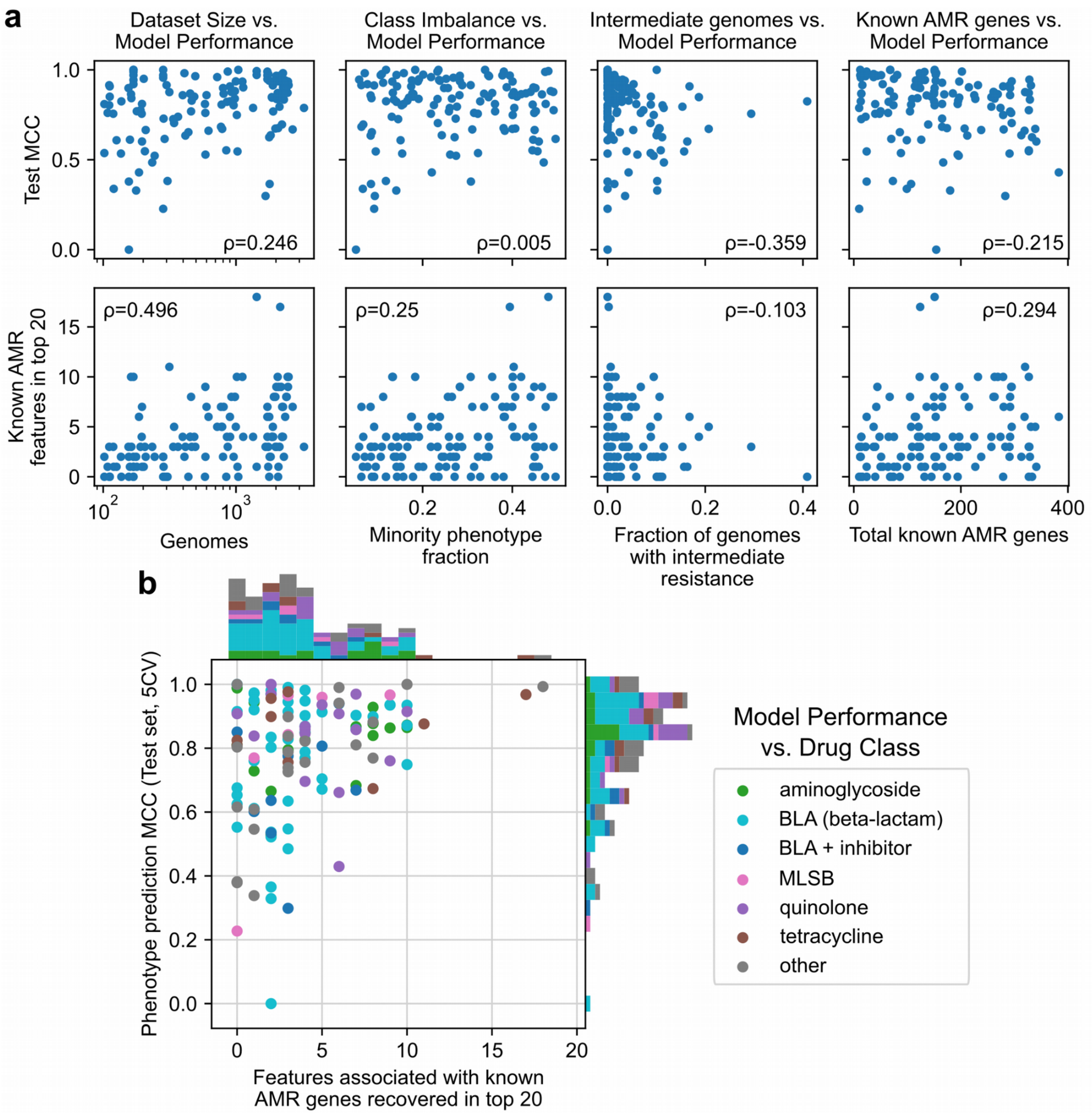


**Figure S10: Relationship between dataset parameters and model performance across 127 species-drug cases.** a) Performance of hyperparameter optimized SVM ensembles vs. dataset size, extent of class imbalance, abundance of "intermediate" resistant genomes, and total known AMR genes annotated. Spearman correlation coefficients are shown. d) Performance of the 127 SVM ensembles versus drug class.

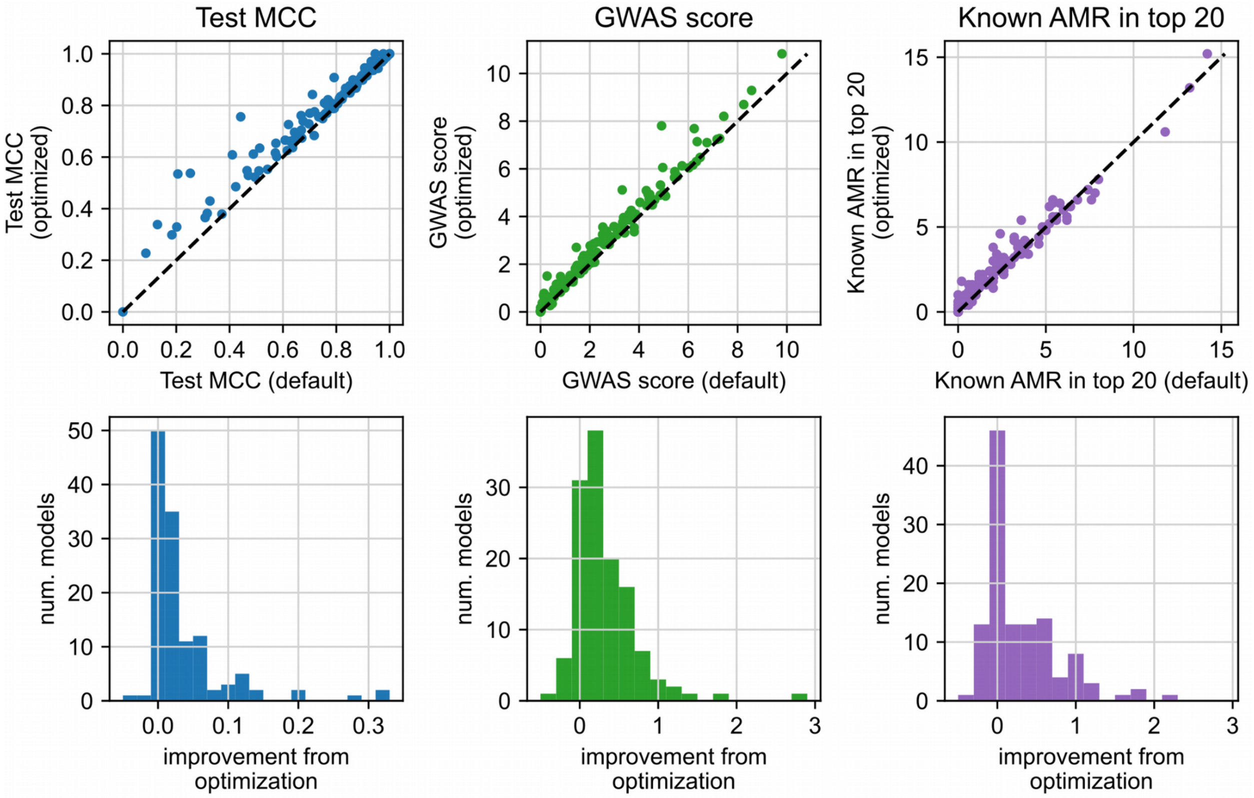


**Figure S11: Performance improvements from hyperparameter optimization of SVM ensembles.** Performance by three metrics is shown for SVM ensembles trained using “default” fixed hyperparameters (C = 1, feature fraction = 50%, sample fraction = 75%, ensemble size = 50) compared to ensembles with hyperparameters optimized to balance both phenotype prediction accuracy and known AMR gene recovery. Performance is based on means from 5-fold cross validation.

**a**

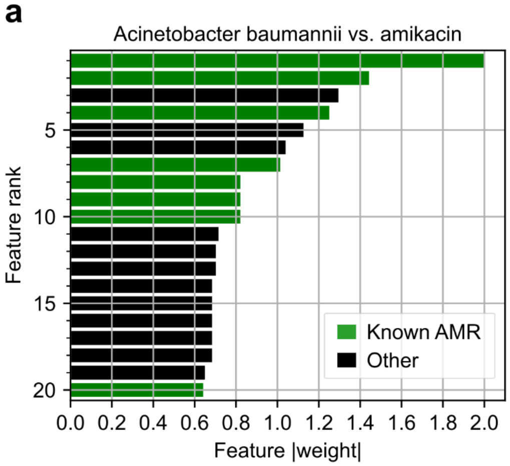


| Rank | Found by Pyseer? | Found by Fisher? | Gene | Feature Type |
|---|---|---|---|---|
| 1 | X | X | APH(3')-VI | gene |
| 2 | - | - | APH(3')-VI | coding_variant |
| 3 | | | hypothetical protein 1 | gene |
| 4 | X | - | armA | gene |
| 5 | | | hypothetical protein 1 | coding_variant |
| 6 | | | hypothetical protein 2 | gene |
| 7 | - | - | APH(3')-VI | coding_variant |
| 8 | - | - | AAC(6')-I | coding_variant |
| 9 | - | - | AAC(6')-I | gene |
| 10 | X | - | armA | coding_variant |
| 11 | | | hypothetical protein 1 | 5_variant |
| 12 | | | mobile element 1 | coding_variant |
| 13 | | | mobile element 1 | gene |
| 14 | | | hypothetical protein 3 | gene |
| 15 | | | hypothetical protein 4 | coding_variant |
| 16 | | | hypothetical protein 3 | coding_variant |
| 17 | | | hypothetical protein 5 | 3_variant |
| 18 | | | hypothetical protein 5 | coding_variant |
| 19 | | | hypothetical protein 1 | 5_variant |
| 20 | X | - | APH(3')-VI | coding_variant |

**Known AMR Features/Genes: 8/3**

**b**

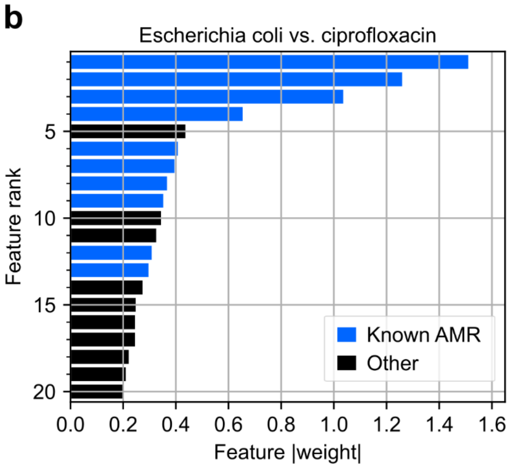


| Rank | Found by Pyseer? | Found by Fisher? | Gene | Feature Type |
|---|---|---|---|---|
| 1 | X | - | gyrA | coding_variant |
| 2 | X | - | parC | coding_variant |
| 3 | - | - | gyrA | coding_variant |
| 4 | X | X | gyrA | coding_variant |
| 5 | | | 5S rRNA | coding variant |
| 6 | - | - | gyrA | coding_variant |
| 7 | X | - | parE | coding_variant |
| 8 | - | - | qnrS | 5_variant |
| 9 | - | - | gyrA | coding_variant |
| 10 | | | hypothetical protein 6 | coding_variant |
| 11 | | | mobile element 2 | coding_variant |
| 12 | - | - | parC | coding_variant |
| 13 | - | - | parC | coding_variant |
| 14 | | | hypothetical protein 7 | 5_variant |
| 15 | | | yeeV | 3_variant |
| 16 | | | eptA | gene |
| 17 | | | eptA | 3_variant |
| 18 | | | yihF | 5_variant |
| 19 | | | parB | 5_variant |
| 20 | | | Mu10 (phage-encoded) | coding_variant |

**Known AMR Features/Genes: 10/4**

**c**

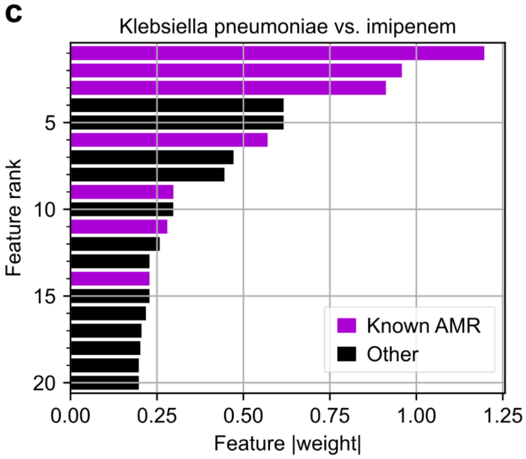


| Rank | Found by Pyseer? | Found by Fisher? | Gene | Feature Type |
|---|---|---|---|---|
| 1 | X | X | blaKPC | gene |
| 2 | X | X | blaKPC | 3_variant |
| 3 | - | - | blaOXA | gene |
| 4 | | | hypothetical protein 8 | coding_variant |
| 5 | | | hypothetical protein 8 | gene |
| 6 | - | - | blaNDM | gene |
| 7 | | | mobile element 3 | gene |
| 8 | | | yeaY | gene |
| 9 | - | - | blaOXA | 3_variant |
| 10 | | | ompC | gene |
| 11 | - | - | blaNDM | coding_variant |
| 12 | | | yafQ | coding_variant |
| 13 | | | hypothetical protein 8 | 3_variant |
| 14 | - | - | blaNDM | 3_variant |
| 15 | | | hypothetical protein 8 | 5_variant |
| 16 | | | srlE | gene |
| 17 | | | hypothetical protein 9 | gene |
| 18 | | | ompC | 5_variant |
| 19 | | | hypothetical protein 10 | coding_variant |
| 20 | | | incW | 5_variant |

**Known AMR Features/Genes: 7/3**

**Figure S12: Visualization of the top 20 genetic features associated with AMR for three species-drug cases as identified by SVM ensembles.** For each feature, the absolute value of the feature weight, overall rank, gene name, and feature type are shown. Features related to known AMR genes are colored and also labeled by whether they were also recovered by Pyseer and/or Fisher's exact test, and other features are black. Features related to undercharacterized genes are labeled as either "hypothetical protein" or "mobile element", numbered by the order the gene is displayed. Additional details are in Dataset S5.

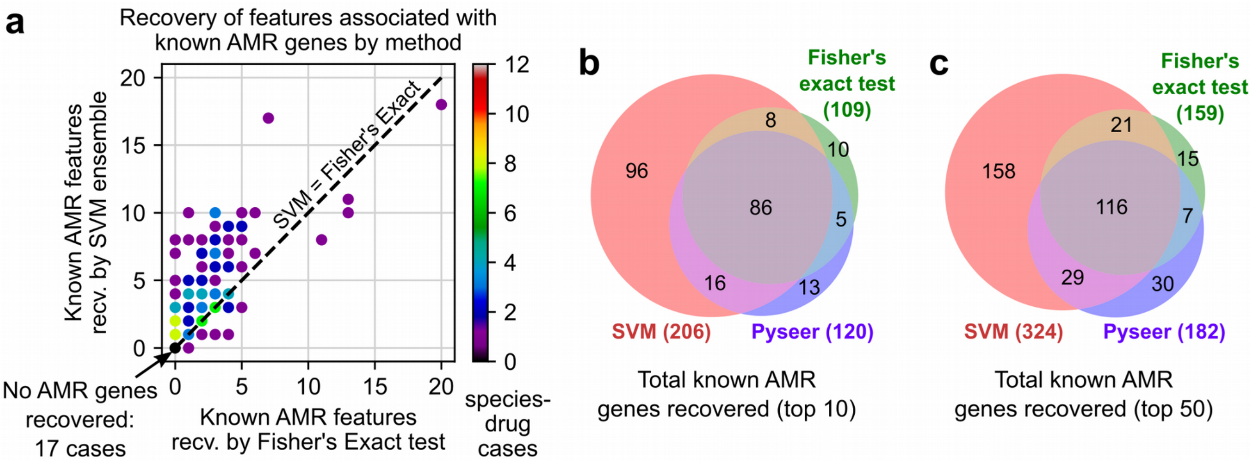


**Figure S13: Additional comparisons between performance of SVM ensembles, Pyseer, and Fisher's exact test at recovering known AMR genes.** a) Comparison between SVMs and Fisher's exact test at recovering known AMR genes across 127 species-drug cases. b-c) Total known AMR gene-drug mappings recovered by SVMs, Pyseer, and Fisher's exact tests across all cases, when defining a recovered gene as those in the top 10 or top 50 features when sorting by feature weight (for SVM) or p-value (for Pyseer and Fisher's exact test).

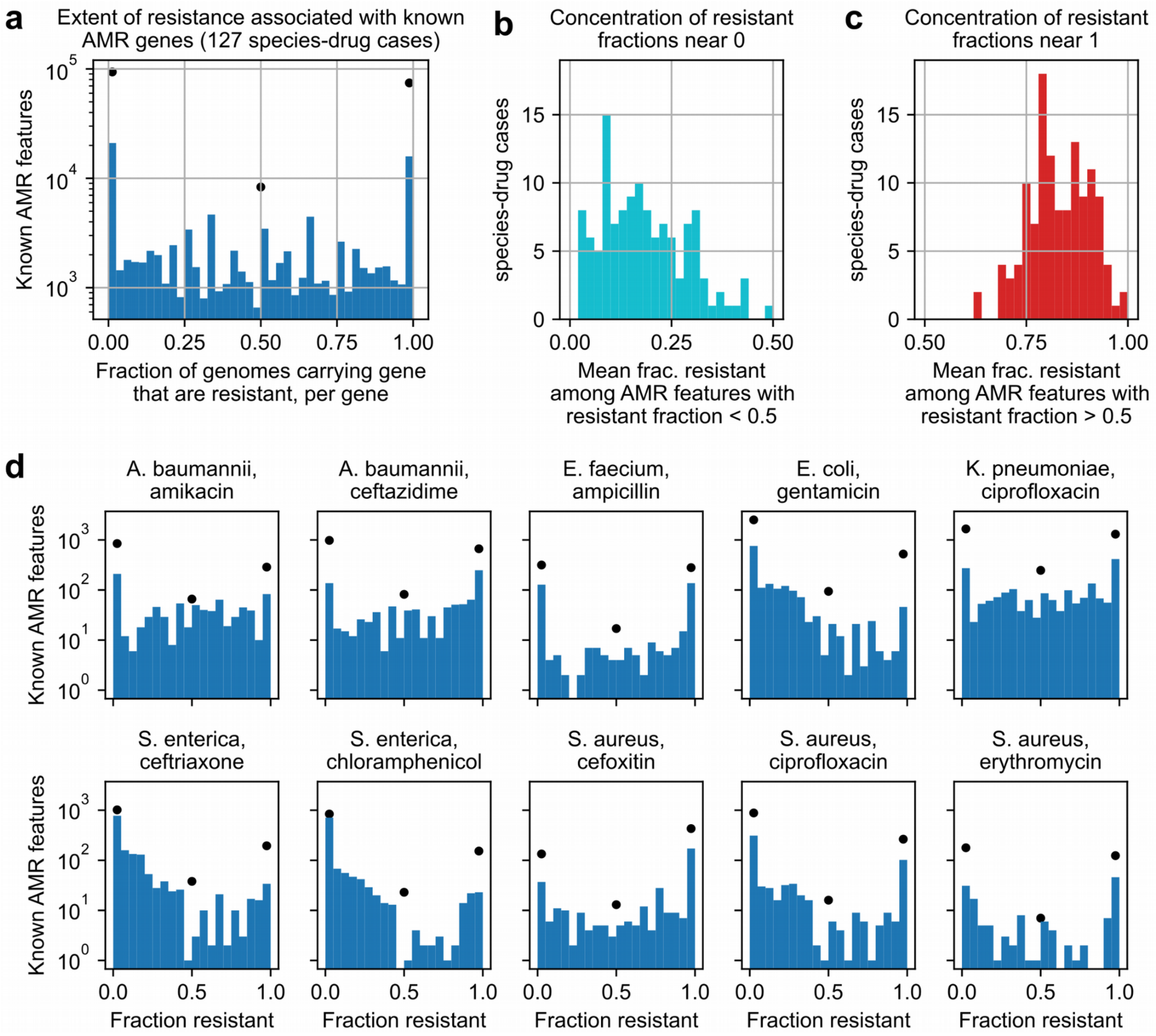


**Figure S14: Associations between known AMR features and resistance.** For each known AMR feature in a given species-drug case, the fraction of genomes with the feature that are resistant to the drug was computed. The distribution of these resistance fractions combined across 127 species-drug cases is shown in (a), with black dots representing rare features (found in no more than 2 genomes) and blue bars representing all other known AMR features. The tendency for these distributions to concentrate near 0 and 1 was quantified by computed means for values b) less than or c) greater than 0.5. Individual distributions for 10 species-drug cases are shown in (d).

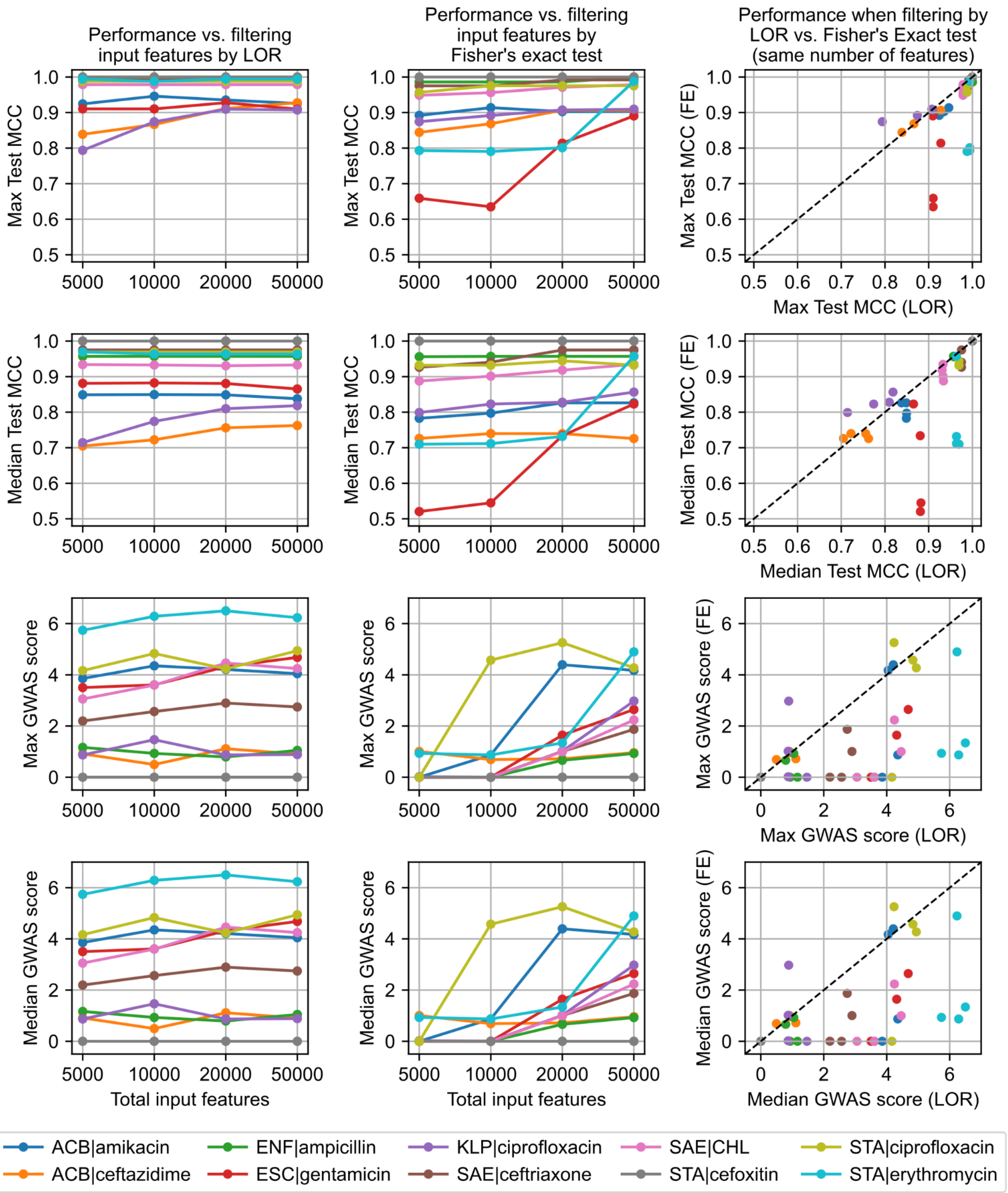


**Figure S15: Impact of varying the preliminary feature filter on downstream model performance.** Each row shows the effect of the feature filter on a model performance metric, either the maximum or median test set MCC (from 5-fold CV) or GWAS score across 256 hyperparameter combinations. The first column shows model performance when limited to the top 5,000, 10,000, 20,000, or 50,000 features after sorting by log odds ratio (LOR) for resistance, and the second column shows model performance under analogous filters sorting instead by Fisher's exact test p-value for resistance. The third column compares performance between LOR and Fisher's exact test filters for identical feature count limits. Each color represents results for a specific species-drug case.

### Supplemental Tables

**Table S1: Genetic feature counts by species.**

| | | ORF-associated | | | | Non-ORF-associated | |
|---|---|---|---|---|---|---|---|
| species | genomes | genes | alleles | 5' flanking variants | 3' flanking variants | noncoding clusters | noncoding variants |
| *A. baumannii* | 1327 | 43888 | 239729 | 262420 | 275216 | 177 | 1725 |
| *C. coli* | 283 | 8299 | 44804 | 32011 | 31527 | 77 | 231 |
| *C. jejuni* | 452 | 9801 | 48710 | 35975 | 36276 | 82 | 381 |
| *E. cloacae* | 484 | 51969 | 325374 | 394577 | 406576 | 167 | 1716 |
| *E. faecium* | 1432 | 53518 | 174176 | 116209 | 113574 | 119 | 890 |
| *E. coli* | 3856 | 148886 | 1024333 | 1005186 | 1003017 | 435 | 5562 |
| *K. pneumoniae* | 3022 | 94147 | 562721 | 584114 | 596130 | 356 | 5010 |
| *N. gonorrhoeae* | 5962 | 18225 | 199856 | 122707 | 117320 | 74 | 693 |
| *P. aeruginosa* | 1050 | 67217 | 448882 | 558844 | 581936 | 136 | 903 |
| *S. enterica* | 3302 | 56315 | 262555 | 255349 | 261757 | 322 | 4932 |
| *S. aureus* | 2248 | 22649 | 210749 | 177169 | 173244 | 135 | 1826 |
| *S. pneumoniae* | 3737 | 36070 | 295716 | 269652 | 281790 | 110 | 1669 |

**Table S2: Enrichment of AMR gene categories in multispecies over single species genes and plasmid over chromosomal genes.** For each comparison, "genes" is the total number of genes in the category, "LOR" is log2 odds ratio, and "p-value" is Fisher's exact test p-value. Starred values are significant at FWER < 0.05 (Bonferroni correction, 36 tests). Categories are sorted by LOR for plasmid over chromosomal genes.

| | **Multispecies over Single species** | | | **Plasmid over Chromosomal** | | |
|---|---|---|---|---|---|---|
| **AMR gene category** | **genes** | **LOR** | **p-value** | **genes** | **LOR** | **p-value** |
| aminoglycoside modifying enzymes (AMEs) | 563 | 1.272 | <0.00001* | 455 | 2.957 | <0.00001* |
| ribosomal protection proteins | 154 | 0.113 | 0.72879 | 115 | 2.348 | <0.00001* |
| chloramphenicol acetyltransferases (CATs) | 91 | 1.001 | 0.00656 | 77 | 1.850 | <0.00001* |
| dihydrofolate reductases (DHFRs), dihydropteroate synthases (DHPSs) | 219 | 1.498 | <0.00001* | 191 | 1.845 | <0.00001* |
| *rpoB* variants | 74 | -0.718 | 0.24769 | 60 | 1.540 | 0.00007* |
| rRNA methyltransferases (rRNA MTases) | 152 | 0.061 | 0.81658 | 132 | 1.049 | 0.00008* |
| beta-lactamases | 645 | 0.015 | 0.90646 | 541 | 1.004 | <0.00001* |
| glycopeptide resistance clusters (RCs) | 425 | -0.721 | 0.00217 | 357 | -0.123 | 0.50527 |
| *gyrA/gyrB/parC* variants | 165 | -1.021 | 0.01013 | 141 | -0.979 | 0.00174 |
| non-two-component system (non-TCS) regulators | 429 | 0.358 | 0.06540 | 388 | -0.993 | <0.00001* |
| penicillin-binding proteins (PBPs) | 254 | -0.257 | 0.41414 | 211 | -1.088 | 0.00002* |
| aminoacyl-tRNA synthetases (AARSs) | 101 | -1.454 | 0.01001 | 82 | -1.094 | 0.00939 |
| efflux | 1519 | -0.215 | 0.08753 | 1346 | -1.107 | <0.00001* |
| phosphoethanolamine (pEtN) transferases | 358 | -1.373 | <0.00001* | 316 | -1.485 | <0.00001* |
| other | 410 | -0.218 | 0.34748 | 362 | -1.811 | <0.00001* |
| two-component system (TCS) regulators | 494 | -0.900 | 0.00005* | 429 | -1.988 | <0.00001* |
| *cya* variants | 57 | -0.544 | 0.45547 | 50 | -3.273 | 0.00002* |
| porins | 74 | -1.595 | 0.01988 | 69 | -3.762 | <0.00001* |

**Table S3: Distribution of the most frequently observed TEM-family beta-lactamases by species.** Mutations are shown relative to the TEM-1 variant. Variants are ordered by total count. Species are abbreviated as *A. baumannii* (AcB), *E. cloacae* (EnC), *E. coli* (EsC), *K. pneumoniae* (KlP), *N. gonorrhoeae* (NeG), *P. aeruginosa* (PsA), *S. enterica* (SaE), and *S. aureus* (StA).

| | Number of genomes with variant by species | | | | | | | | | |
|---|---|---|---|---|---|---|---|---|---|---|
| **Mutations** | **Variant** | **AcB** | **EnC** | **EsC** | **KlP** | **NeG** | **PsA** | **SaE** | **StA** | **Total** |
| - | TEM-1 | 414 | 171 | 1692 | 1179 | 235 | 1 | 732 | - | 4424 |
| M180T | TEM-135 | - | - | 8 | 1 | 147 | - | 2 | - | 158 |
| P12S | * | - | - | - | - | 108 | - | - | - | 108 |
| V82I, A182V | TEM-116 | - | - | - | - | - | - | 17 | 11 | 28 |
| M67I | TEM-40 | - | - | 16 | 1 | - | - | - | - | 17 |
| R241S | TEM-30 | - | - | 16 | - | - | - | - | - | 16 |
| Q37K | TEM-2 | - | 14 | - | 1 | - | - | - | - | 15 |
| M67L | TEM-33 | - | - | 7 | 1 | - | - | 2 | - | 10 |
| 14insMQQCL | * | - | - | - | - | - | - | 10 | - | 10 |

* No exact match in the CARD database

**Table S4: Initial test species-drug cases for hyperparameter testing.** Columns “n”, “% Sus.” and “AMR genes” refer to the number of genomes with AMR data, the percent of those genomes that are susceptible, and the number of known AMR genes identified that are associated with the drug for that species, respectively.

| Species | Species Class | Drug | Drug Class | n | % Sus. | AMR genes |
|---|---|---|---|---|---|---|
| ***A. baumannii*** | γ-proteobacteria | **amikacin** | aminoglycoside | 924 | 49.1 | 150 |
| ***A. baumannii*** | γ-proteobacteria | **ceftazidime** | beta-lactam | 960 | 15.3 | 151 |
| ***E. coli*** | γ-proteobacteria | **gentamicin** | aminoglycoside | 2447 | 86.6 | 263 |
| ***K. pneumoniae*** | γ-proteobacteria | **ciprofloxacin** | quinolone | 2089 | 19.1 | 227 |
| ***S. enterica*** | γ-proteobacteria | **ceftriaxone** | beta-lactam | 2259 | 84.2 | 154 |
| ***S. enterica*** | γ-proteobacteria | **chloramphenicol** | other | 2348 | 82.4 | 116 |
| ***E. faecium*** | Bacilli | **ampicillin** | beta-lactam | 1432 | 14.2 | 81 |
| ***S. aureus*** | Bacilli | **cefoxitin** | beta-lactam | 1041 | 23.5 | 68 |
| ***S. aureus*** | Bacilli | **ciprofloxacin** | quinolone | 1725 | 62.2 | 42 |
| ***S. aureus*** | Bacilli | **erythromycin** | other | 2014 | 71.7 | 44 |

**Table S5: Negative log10 p-values for Kruskal-Wallis tests between SVM ensemble hyperparameters and model performance.** Performance metrics compared are phenotype prediction performance (mean test set MCC during 5-fold cross validation) and recovery of known AMR genes among model features (GWAS score).

| | Hyperparameter vs. Test set MCC | | | | Hyperparameter vs. GWAS score | | | |
|---|---|---|---|---|---|---|---|---|
| AMR Case | SVM C | feature fraction | sample fraction | ensemble size | SVM C | feature fraction | sample fraction | ensemble size |
| *A. baumannii,* amikacin | 1.009 | 12.123 | 6.016 | 0.75 | 1.967 | 26.219 | 17.297 | 0.711 |
| *A. baumannii,* ceftazidime | 17.258 | 0.187 | 24.916 | 0.58 | 3.212 | 0.402 | 39.543 | 0.011 |
| *E. coli,* gentamicin | 3.256 | 10.935 | 31.611 | 0.051 | 2.963 | 35.68 | 1.47 | 5.058 |
| *E. faecium,* ampicillin | 8.738 | 11.894 | 17.011 | 0.307 | 7.905 | 22.447 | 9.831 | 1.674 |
| *K. pneumoniae,* ciprofloxacin | 20.421 | 1.621 | 19.877 | 0.051 | 28.08 | 4.177 | 2.767 | 3.495 |
| *S. aureus,* cefoxitin | 48.629 | 0.117 | 0.353 | 0.001 | 14.449 | 10.696 | 1.244 | 0.001 |
| *S. aureus,* ciprofloxacin | 9.228 | 21.314 | 10.536 | 0.416 | 1.211 | 26.275 | 18.174 | 0.059 |
| *S. aureus,* erythromycin | 3.635 | 10.07 | 22.577 | 0.723 | 3.065 | 40.817 | 6.849 | 0.158 |
| *S. enterica,* ceftriaxone | 0.791 | 4.16 | 36.582 | 0.618 | 4.264 | 42.314 | 1.036 | 0.59 |
| *S. enterica,* chloramphenicol | 6.62 | 0.579 | 30.051 | 1.804 | 5.181 | 14.501 | 3.774 | 2.055 |

**Table S6: SVM hyperparameter ranges used during optimization.** Larger initial ranges were used for evaluating the 10 test species-drug cases, from which reduced ranges were derived for evaluating against all 127 species-drug cases.

| Hyperparameter | Initial range | Reduced range |
|---|---|---|
| Number of estimators | 25, 50, 100, 200 | 25, 50 |
| Fraction of samples per estimator | 25%, 50%, 75%, 100% | 50%, 75%, 100% |
| Fraction of features per estimator | 25%, 50%, 75%, 100% | 25%, 50%, 75%, 100% |
| C (SVM regularization term) | 0.1, 1, 10, 100 | 0.1, 1, 10 |
| **Unique combinations** | 256 | 72 |

**Table S7: Statistically significant differences in model performance between selected and excluded hyperparameter combinations for 10 species-drug cases.** For each case, a Mann-Whitney U-test was conducted to determine if the trained SVM model's test set MCCs or GWAS scores, across all hyperparameter combinations (HPCs) and folds from 5-fold CV, differed significantly for selected HPCs than for excluded HPCs. See Table S6 for HPC sets. Starred cases are significant to FWER < 0.05, Bonferroni correction (20 tests).

| | **Test set MCC** | | | **GWAS Score** | | |
|---|---|---|---|---|---|---|
| **AMR Case** | Included Median | Excluded Median | MW test p-value | Included Median | Excluded Median | MW test p-value |
| *A. baumannii*, amikacin | 0.83782 | 0.83815 | 0.23016 | 4.04567 | 4.07742 | 0.04913 |
| *A. baumannii*, ceftazidime | 0.76253 | 0.75629 | 0.00138* | 0.9075 | 0.59558 | <0.00001* |
| *E. faecium*, ampicillin | 0.95698 | 0.95698 | 0.12555 | 1.04808 | 1.07578 | 0.1873 |
| *E. coli*, gentamicin | 0.86522 | 0.86143 | <0.00001* | 4.68256 | 5.1806 | <0.00001* |
| *K. pneumoniae*, ciprofloxacin | 0.81835 | 0.81573 | 0.41189 | 0.88562 | 0.87238 | 0.34162 |
| *S. enterica*, ceftriaxone | 0.9751 | 0.9751 | 0.04874 | 2.74439 | 3.12598 | <0.00001* |
| *S. enterica*, chloramphenicol | 0.93276 | 0.93276 | 0.12058 | 4.24349 | 4.28149 | 0.77218 |
| *S. aureus*, cefoxitin | 1.00000 | 1.00000 | 0.94434 | 0.00000 | 0.00000 | <0.00001* |
| *S. aureus*, ciprofloxacin | 0.96917 | 0.96917 | 0.60248 | 4.94068 | 4.67576 | 0.00018* |
| *S. aureus*, erythromycin | 0.96318 | 0.9575 | 0.04671 | 6.23253 | 6.23332 | 0.37441 |

**Table S8: Impact of input data properties on SVM ensemble performance.** Impact was assessed with Kruskal-Wallis tests for categorical properties (species, drug class) and Spearman correlation for quantitative properties (number of genomes, minority phenotype fraction, fraction of genomes with the "intermediate" phenotype, total number of known AMR genes). Rows are sorted by p-value. Cases significant to FWER < 0.05 (Bonferroni correction, 12 tests) are starred.

| Performance Metric | Data Metric | Statistical Test | p-value |
|---|---|---|---|
| AMR genes in top 20 | num. genomes | Spearman-R | <0.00001* |
| Test MCC in 5CV | species | Kruskal-Wallis | <0.00001* |
| Test MCC in 5CV | intermediate fraction | Spearman-R | 0.00003* |
| AMR genes in top 20 | total known AMR genes | Spearman-R | 0.00079* |
| AMR genes in top 20 | minority fraction | Spearman-R | 0.00464 |
| Test MCC in 5CV | num. genomes | Spearman-R | 0.00537 |
| AMR genes in top 20 | species | Kruskal-Wallis | 0.00721 |
| Test MCC in 5CV | total known AMR genes | Spearman-R | 0.01536 |
| AMR genes in top 20 | intermediate fraction | Spearman-R | 0.25039 |
| Test MCC in 5CV | drug class | Kruskal-Wallis | 0.30250 |
| AMR genes in top 20 | drug class | Kruskal-Wallis | 0.48732 |
| Test MCC in 5CV | minority fraction | Spearman-R | 0.95993 |

**Table S9: Summary of known AMR gene-drug mappings recovered by Fisher's Exact test but missed by the SVM ensemble approach.** Drugs abbreviated are quinupristin-dalfopristin (Q-D) and trimethoprim-sulfamethoxazole (SXT)

| Species | Drug | Missed Gene | Model Test MCC (5CV) | Gene Rank in Model | Statistically Unique Features in Top 50* | Possible Failure Mode |
|---|---|---|---|---|---|---|
| *Acinetobacter baumannii* | tobramycin | *aadA*** | 0.87 | 33 | 44 | Feature rank below top 20 threshold |
| *Enterobacter cloacae* | cefepime | *blaKPC*** | 0.61 | - | 46 | Poor model performance |
| *Enterococcus faecium* | Q-D | *eatAv* | 1.00 | - | 1 | High correlation among top features |
| *Enterococcus faecium* | teicoplanin | *vanXA* | 1.00 | 21 | 11 | Feature rank below top 20 threshold |
| *Enterococcus faecium* | teicoplanin | *vanA* | 1.00 | 21 | 11 | Feature rank below top 20 threshold |
| *Enterococcus faecium* | teicoplanin | *vanHA* | 1.00 | 21 | 11 | Feature rank below top 20 threshold |
| *Enterococcus faecium* | vancomycin | *vanZA*** | 0.99 | - | 16 | High correlation among top features |
| *Enterococcus faecium* | vancomycin | *vanYA*** | 0.99 | - | 16 | High correlation among top features |
| *Escherichia coli* | norfloxacin | *parC* | 0.43 | - | 36 | Poor model performance |
| *Klebsiella pneumoniae* | SXT | *sul1* | 0.84 | - | 13 | High correlation among top features |
| *Neisseria gonorrhoeae* | erythromycin | *mtrR* | 0.77 | - | 48 | Poor model performance |
| *Staphylococcus aureus* | ciprofloxacin | *arlR* | 0.97 | - | 46 | - |

*Refers to the number of features remaining among the SVM model's top 50 features by weight after collapsing perfectly correlated features together. Lower values correspond to more highly correlated features.

**Also in the top 20 features by Pyseer.

**Table S10: 13 gene flanking noncoding variants predicted to be associated with resistance against specific drug classes for individual species.** Accession IDs are provided for the most common allele of the corresponding gene cluster (RefSeq when possible, GenBank otherwise), along with gene names when available and gene products. Mutations are defined relative to the most common 5'/3' variant for the corresponding gene. The number of resistant/susceptible genomes and log2 odds ratios (LORs) for resistance are shown for the top three drugs by LOR when data for more than three related drugs was available. Drug abbreviations and mappings to relevant sequences are available in Dataset S7.

| Species | Drug Class | Accession (Gene) | Predicted Gene Product | Mutations* | Resistant/ Susceptible | LORs |
|---|---|---|---|---|---|---|
| | | | *5' flanking variants* | | | |
| *Acinetobacter baumannii* | beta-lactam | AGQ10471.1 (*pqqA*) | Coenzyme PQQ synthesis protein A | -12_2delTGATTTAATCAAGTG** | CTX=73/0<br>CRO=73/0<br>AMP=73/0 | CTX=6.4<br>CRO=6.4<br>AMP=6.4 |
| *Acinetobacter baumannii* | tetracycline | WP_000096554.1 (*clpV*) | T6SS AAA+ chaperone | Most common variant | TET=241/12<br>MIN=71/18 | TET=3.3<br>MIN=2.3 |
| *Campylobacter coli* | quinolone | WP_002805020.1 | GNAT acetyl-transferase | Most common variant | CIP=94/71<br>NAL=95/70 | CIP=6.0<br>NAL=5.5 |
| *Campylobacter coli* | quinolone | WP_002783313.1 | Putative transmembrane transport protein, MFS | Most common variant | NAL=97/115<br>CIP=95/117 | NAL=5.5<br>CIP=5.4 |
| *Escherichia coli* | beta-lactam | AAG56074.1 (*ymgF*) | Small inner membrane protein | -19A>C, -23A>G, -25_24delTC, -30insA, -48G>A, -173A>T, -175C>A, -211A>T, -213C>T | CTX=28/0<br>CAZ=30/1<br>CXM=14/1 | CTX=6.7<br>CAZ=5.7<br>CXM=4.6 |
| *Escherichia coli* | quinolone | WP_000017703.1 (*hybB*) | Ni/Fe-hydrogenase 2 b-type cytochrome subunit | Most common variant | LVX=260/56<br>CIP=606/606<br>NAL=45/16 | LVX=3.3<br>CIP=2.8<br>NAL=1.4 |
| *Streptococcus pneumoniae* | beta-lactam | WP_000449822.1 | Thiaminase II | Most common variant | AMX=13/4<br>MEM=16/1<br>CXM=16/1 | AMX=12.4<br>MEM=9.6<br>CXM=9.3 |
| *Streptococcus pneumoniae* | beta-lactam | ADI69655.1 (*nplT*) | Neopullulanase | Most common variant | AMX=13/3<br>MEM=15/1<br>CXM=15/1 | AMX=12.7<br>MEM=9.1<br>CXM=8.6 |
| *Streptococcus pneumoniae* | beta-lactam | WP_000592948.1 (*ugl*) | Unsaturated chondroitin disaccharide hydrolase | -186C>T | AMX=13/6<br>CXM=17/2<br>MEM=17/3 | AMX=11.9<br>CXM=9.6<br>MEM=9.2 |
| | | | *3' flanking variants* | | | |
| *Campylobacter coli* | quinolone | WP_002777456.1 (*hspR*) | Transcriptional repressor of DnaK operon | Most common variant | NAL=98/115<br>CIP=96/117 | NAL=7.4<br>CIP=7.4 |

| | | | | | | |
|---|---|---|---|---|---|---|
| *Klebsiella pneumoniae* | beta-lactam | WP_000679427.1 (*qacEΔ1*) | Small multidrug resistance (SMR) efflux transporter | Most common variant | AMP=759/0<br>CEF=27/0<br>CRO=772/4 | AMP=10.3<br>CEF=5.5<br>CRO=4.3 |
| *Salmonella enterica* | quinolone | WP_012772747.1 | Psp operon transcriptional activator | Most common variant | NAL=9/11<br>CIP=9/10 | NAL=3.5<br>CIP=3.2 |
| *Streptococcus pneumoniae* | beta-lactam | WP_000145597.1 (*nplT*) | Neopullulanase | Most common variant | AMX=13/3<br>MEM=15/1<br>CXM=15/1 | AMX=12.7<br>MEM=9.1<br>CXM=8.6 |

*Mutations are denoted relative to the start codon for 5' variants. Position -1 corresponds to the first base pair immediately adjacent on the 5' side of the gene's start codon.

**Results in the deletion of a GTG start codon and the 12 base pairs immediately adjacent on the 5' side, with respect to the most common variant. The candidate variant begins with an ATG start codon.

### Supplemental Dataset Captions

**DatasetS1.json: PATRIC genome IDs for all genomes used.**

**DatasetS2.xlsx: Consolidated SIR phenotypes derived from directly reported SIRs and inference from MICs available on PATRIC.** Also includes genome MICs, MIC-SIR mappings used for SIR inference, most common testing standard for each species-drug case determined from PATRIC annotations or manual curation of contributing BioProjects, and distributions of genome MLST subtypes and BioProjects.

**DatasetS3.xlsx: Distribution of unique AMR genes and cross-species AMR gene analysis.** Counts of gene-drug mappings are provided for each species-drug case, along with AMR gene category abbreviations, drug class assignments, curation of AMR gene annotations from PATRIC, and assignments of AMR genes to specific drugs. Species distributions, localization predictions, and function classifications for re-clustered cross-species AMR genes are also included.

**DatasetS4.xlsx: Distribution of complete blaTEM alleles detected and plasmid predictions for contigs containing TEM-116.** Also includes MASH distances between all contigs in genomes containing TEM-116 and PLSDB reference plasmids.

**DatasetS5.xlsx: Summary of SVM model performance and top predictive features across 127 species-drug cases.** Includes dataset properties (number of susceptible/resistant genomes, known AMR genes identified), SVM mean test set MCC across 5-fold cross validation, final hyperparameter choices, known AMR genes recovered by either SVM, Pyseer, or Fisher’s exact test, and raw lists of top 50 features for each model. Also includes data used in the analysis of individual recovered *gyrA* alleles, comparison between preliminary input feature filters, and GWAS score generalizability analysis.

**DatasetS6.zip: Sequences associated with the top 50 features from each SVM model.** Exact sequences for all such features are provided, as well as the top two most common variants of each type for all sequence clusters related to the features.

**DatasetS7.xlsx: Filtering results for identifying and categorizing novel AMR gene candidates from SVM models.** Also includes abbreviations for drug names.

**DatasetS8.xlsx: Cell densities achieved by *cycA* and *frdD* mutants under various antibiotic stresses, base media, and supplements.** Results of statistical tests between densities achieved for different strains or conditions and predicted *ampC* transcription rates for *frdD* mutants are also included. Also includes all instances of *cycA*, *frdD*, and *ampC* identified for all *E. coli* genomes in this study.
